# An electrostatic relay connects client binding and liquid-liquid phase separation of tau

**DOI:** 10.64898/2026.01.21.700558

**Authors:** Hannah Osterholz, Gefei Chen, Jakob Freudenberger, Cecilia Mörman, Axel Abelein, Dilraj Lama, Michael Landreh, Axel Leppert

## Abstract

Intrinsically disordered proteins (IDPs) achieve functional specificity through dynamic, multivalent interactions that are often reorganized during liquid–liquid phase separation (LLPS). How LLPS reshapes IDP conformational landscapes and binding preferences remains poorly understood. Here, we combine native ion mobility mass spectrometry, mass photometry, fluorescence microscopy, and computational modeling to dissect how conditions that promote LLPS of the intrinsically disordered human tau protein regulate its interactions with tubulin and the neuronal anti-amyloid chaperone domain BRICHOS from Bri2. We show that electrostatically driven compaction and transient oligomerization of tau during LLPS promote BRICHOS binding to a specific Tau segment. Within tau condensates, BRICHOS can compete with tubulin for tau interactions by blocking its adjacent binding site, thereby modulating LLPS-dependent microtubule assembly. Using a generalizable experimental strategy, we provide a proof of concept for detecting conformational selectivity in a dynamic condensate.

## Introduction

Intrinsically disordered proteins (IDPs) are central to cellular regulation and function, yet their lack of defined secondary and tertiary structures raises fundamental questions about how their interactions are regulated. Unlike folded proteins, IDPs populate dynamic conformational ensembles and often engage with binding partners through weak, multivalent interactions ^1,2^. Some IDPs contain well-defined interaction sites that fold upon binding to their partners, whereas others rely on dispersed sequence features, such as clusters of charged or aromatic residues, that mediate low-affinity contacts ^3–5^. These multivalent interactions are also central to liquid-liquid phase separation (LLPS) of IDPs, where they mediate demixing of the proteins into dense condensates ^3^. These liquid-like, membraneless condensates typically form via π–π, cation–π, dipolar, and electrostatic contacts between low-complexity (LC) sequences. The short-lived nature of these interactions ensures that condensates remain dynamic and in constant exchange with their environment ^6–8^. By organizing biomolecules into highly concentrated microenvironments, LLPS enables IDPs to act as hubs in cell signaling, transcription, translation, and stress responses ^8,9^. However, many of the underlying interactions are transient and experimentally difficult to study, leaving key aspects such as the regulation of IDP-partner engagement unresolved.

A prominent example of an intrinsically disordered protein is tau, whose ability to undergo LLPS has been implicated in regulating aspects of its cellular function ^10^. Full-length tau (termed 2N4R) is composed of an N-terminal region, a proline-rich domain (PRD), a microtubule-binding domain (MTBD), and a C-terminal region, all of which are largely disordered with a high predicted propensity to undergo LLPS (Figure S1a) ^11^. The tau sequence contains multiple clusters of charged residues that promote multivalent interactions important for LLPS (Figure S1b) ^12^. Importantly, phase-separated tau sequesters and concentrates tubulin, and in this manner promotes microtubule polymerization ^13^. Tau also binds along the microtubule lattice to stabilize microtubules and regulate cytoskeletal dynamics ^14,15^. However, under pathological conditions, tau can misassemble into amyloid fibrils that accumulate into neurofibrillary tangles associated with several neurodegenerative disorders ^16,17^. LLPS has emerged as a potential intermediate along this pathogenic pathway, where aggregation-prone regions of tau become locally concentrated, facilitating aberrant self-assembly ^18^. Therefore, cells tightly regulate multivalent interactions in condensates through selective recruitment of various modulators that reshape conformational ensembles ^19,20^. For tau, molecular chaperones, post-translational modifications, RNA, and small molecules have all been shown to modulate its macroscopic phase behavior and its propensity to adopt aggregation-prone states ^21–23^.

In this work, we dissect the molecular determinants shaping interaction profiles within tau condensates. To overcome the experimental challenges posed by transient, multivalent, and LLPS-dependent interactions, we combine native mass spectrometry, mass photometry, and fluorescence microscopy with computational analyses to resolve how tau engages different binding partners under conditions that promote the dispersed or condensed states. This strategy enables us to define LLPS-dependent binding preferences and uncover interaction regions enabled by LLPS-associated conformational changes. Our integrated approach provides a generalizable framework that reveals how condensate pathways regulate protein-protein interactions.

## Results

### Chaperone recruitment inhibits microtubule assembly in tau condensates

Tau is known to undergo LLPS under low salt conditions, where condensate formation is mediated by intermolecular electrostatic interactions. Tau readily formed condensates at 2.5 mM ammonium acetate (AmAc), a volatile salt suitable for mass spectrometric analysis of proteins (Figure 1a). We then assessed client protein recruitment into tau condensates. For this purpose, fluorescently labeled protein was added to fluorescently labeled tau under droplet promoting conditions and co-localization was assessed by fluorescence microscopy. As expected, tubulin was readily recruited into tau droplets formed in 2.5 mM AmAc. Next, we tested recruitment of the Bri2 BRICHOS domain from the Bri2 protein (integral membrane protein 2B), an anti-amyloid chaperone that incorporates into tau condensates and modulates condensate dynamics ^21^. In addition, Bri2 BRICHOS associates with the neuronal cytoskeleton *in vivo* and *in vitro* ^24,25^. Like tubulin, the Bri2 BRICHOS variant R221E, a monomeric variant with increased anti-amyloid chaperone capacity (hereafter referred to as BRICHOS) ^26^, co-localized with tau in droplets (Figure 1a), demonstrating that tubulin and BRICHOS are recruited into condensates under identical conditions.

**Figure 1.**
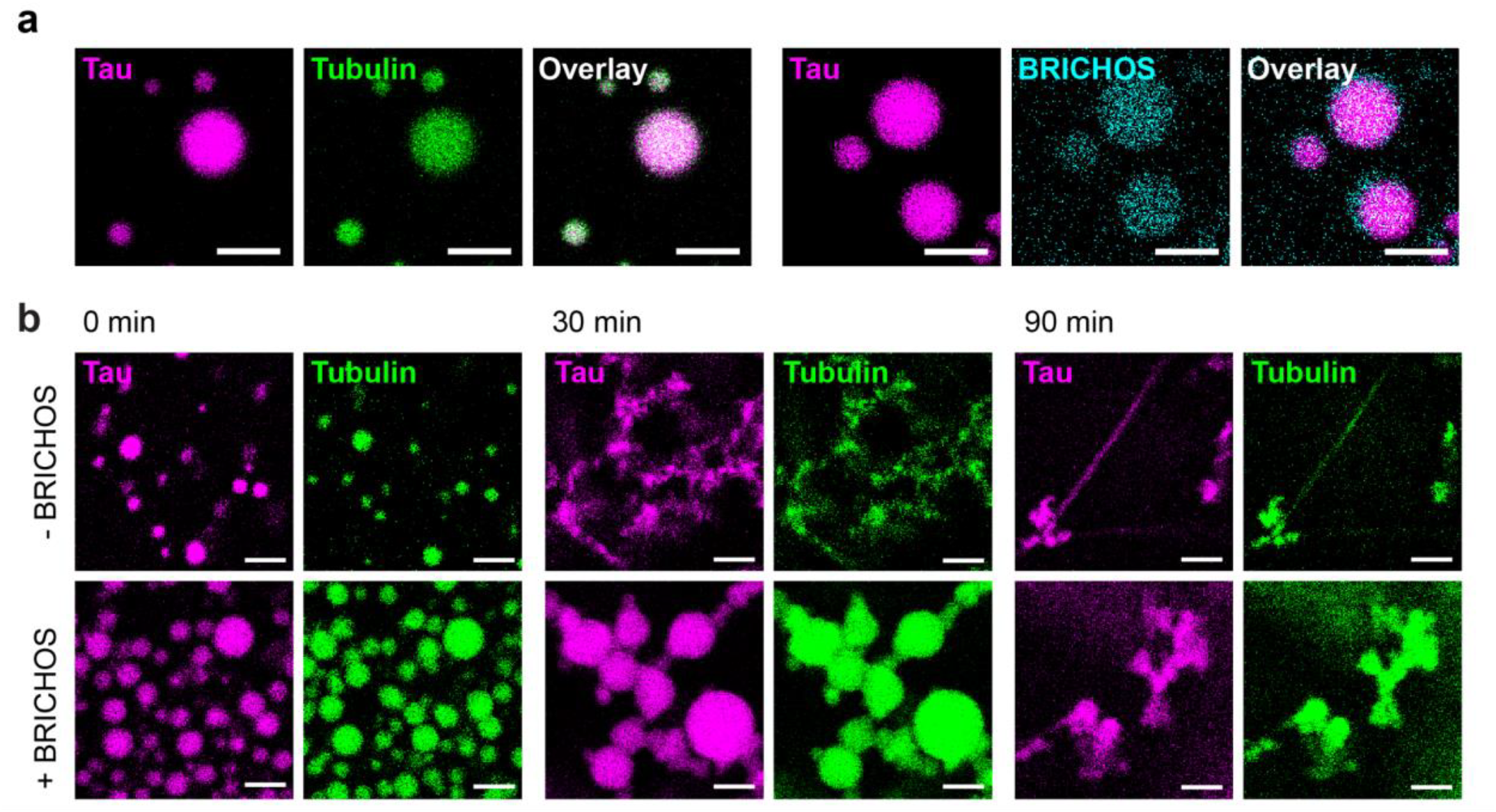
BRICHOS is recruited into tau condensates and impairs tubulin assembly. (a) Fluorescence microscopy of 20 µM tau (+ 1% tau-atto655), 0.4 µM tubulin-atto488 and 0.8 µM BRICHOS-atto390 in 2.5 mM AmAc pH 8 show colocalization in tau droplets at room temperature. Scale bars, 5 μm. (b) Fluorescence microscopy of 20 µM tau (+2% tau-atto655) and 5 µM tubulin (+5% tubulin-atto488) with 2 µM BRICHOS in 2.5 mM AmAc pH 8. Tau and tubulin colocalize in droplets (0 min). In absence of BRICHOS and after addition of 1 mM GTP, droplets lose their shape (30 min) and microtubule growth is observed (90 min). In the presence of BRICHOS, droplets initially exhibit continued Ostwald maturation (30 min) before eventually converting to non-fibrillar aggregates (90 min). Samples have been incubated at 37°C. Scale bars, 5 μm.

To assess tau-mediated tubulin assembly within condensates, we incubated droplets containing 20 µM tau and 5 µM tubulin at 37°C after addition of 1 mM GTP. We observed that tau droplets began to lose their spherical morphology within 30 min, and within 90 min, microtubule fibrils with co-localized tau extended from the droplets (Figure 1b). Addition of 2 µM BRICHOS together with 5 µM tubulin did not alter the initial droplet appearance. However, over time the droplets increased in size, an effect previously reported upon BRICHOS recruitment into tau condensates ^21^, and instead of fibrillar microtubule-like structures, non-fibrillar tau–tubulin assemblies were observed. Together, these results suggest that BRICHOS recruitment into tau condensates affects tau-mediated tubulin assembly.

### LLPS-promoting conditions induce tau compaction and oligomerization

The inhibitory effect of BRICHOS on tau-mediated tubulin assembly raises the question whether BRICHOS may distort tubulin-tau interactions in condensates. We therefore turned to native ion mobility mass spectrometry (IMMS), which can reveal how LLPS-promoting conditions impact the conformational ensembles and interactions of condensate-forming proteins ^27,28^. In native IMMS, proteins are gently transferred into the gas phase, preserving key features of their solution conformations: native MS data report charge-state distributions and oligomeric states, while ion mobility arrival times inform about the overall shape of individual protein ions. Widefield fluorescence microscopy showed that lowering the AmAc concentration from 100 mM to 12.5 mM led to the onset of droplet formation, while further reduction to 2.5 mM produced abundant condensates that increased in size and number (Figure 2a). With IM-MS, we observed that lower AmAc concentrations resulted in a progressive reduction of the average charge state of tau (Figure 2b and Figure S2b), consistent with an increase in intramolecular interactions.

**Figure 2.**
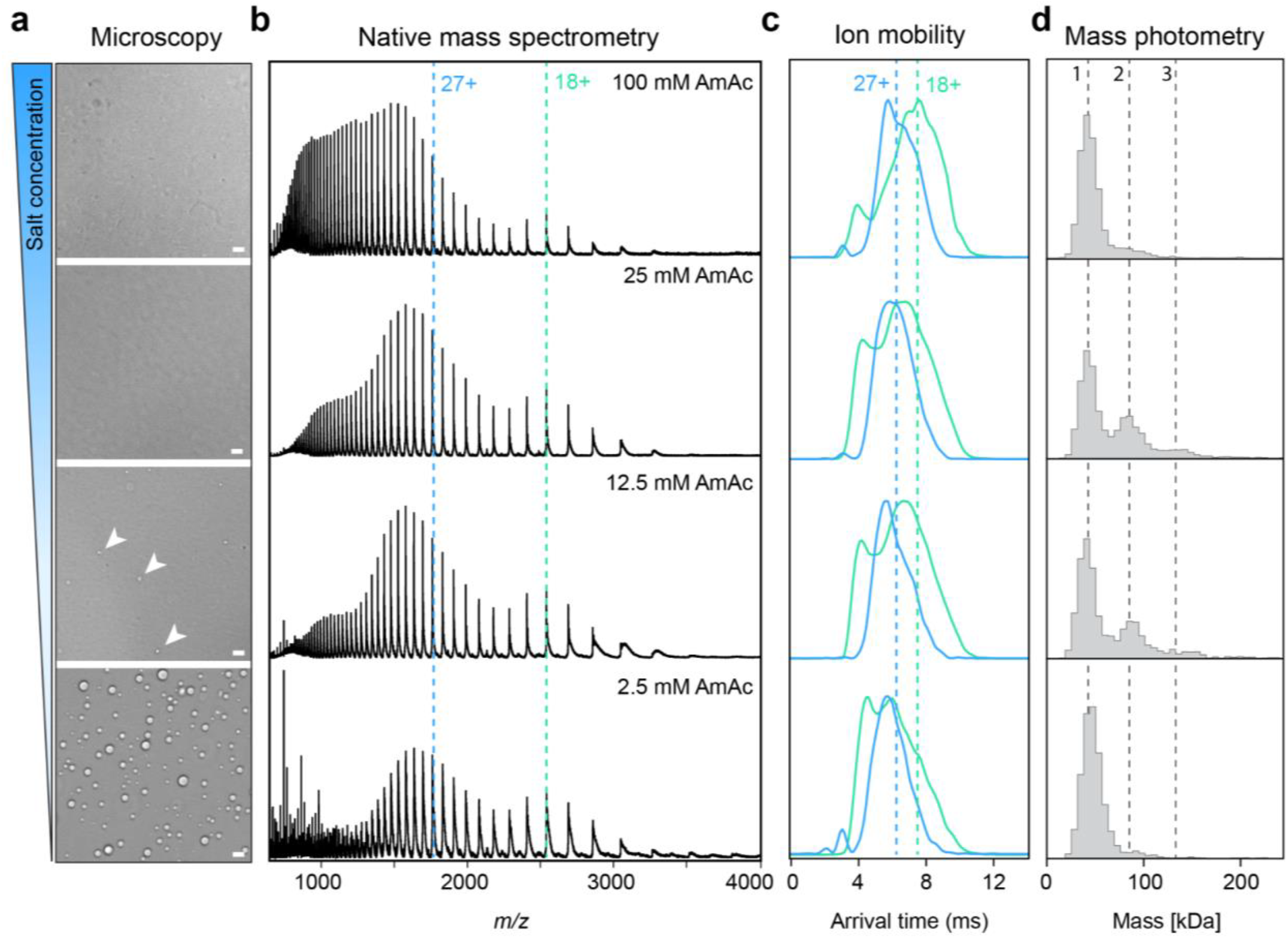
Ion mobility mass spectrometry and mass photometry capture conformational changes and self-association of tau under LLPS-promoting conditions. (a) Widefield microscopy of 20 μM tau in 100 mM, 25 mM, 12.5 mM, 2.5 mM AmAc pH 8 (top to bottom) shows droplet formation at low AmAc concentrations. Scale bars, 5 μm. (b) Native mass spectra of 20 μM tau in 100 mM, 25 mM, 12.5 mM, 2.5 mM AmAc pH 8 (top to bottom) show decreasing average charge states with decreasing salt concentration. (c) Ion mobility of 20 μM tau in 100 mM, 25 mM, 12.5 mM, 2.5 mM AmAc pH 8 (top to bottom) shows compaction of the protein ions with decreasing salt concentration. Dashed lines indicate the center of the arrival time distributions in 100 mM AmAc. (d) MP histograms of 20 nM tau in 100 mM, 25 mM, 12.5 mM, 2.5 mM AmAc pH 8 (top to bottom). At 25 mM and 12.5 mM AmAc tau dimers and tetramers were detected in addition to the monomers visible at all tested concentrations. Dashed lines indicate the molecular weights of tau monomers, dimers, and trimers.

This interpretation agrees with previous work showing that native MS reliably reports on the conformational landscape of proteins with low-complexity domains, where lower charge states reflect more compact conformational ensembles ^29–31^. To exclude progressive solution acidification during ESI as the cause of the reduced charge state at low salt ^32^, we added 1% formic acid to tau in 100 mM AmAc (pH 8). Acidification increased the charge state well beyond any LLPS-promoting condition, demonstrating that the charge-state reduction at low salt arises from tau compaction rather than from an acidification effect (Figure S2a, b). Consistent with this interpretation from the average charge states, IMMS measurements showed a clear decrease in arrival time for identically charged tau ions with decreasing AmAc concentration (Figure 2c). This effect was most pronounced for lower charge states, likely due to strong Coulombic repulsion at high charge states limiting further compaction of the protein ion.

To additionally monitor whether changes in AmAc concentration affect not only intra-but also intermolecular interactions of tau, we turned to mass photometry (MP). Importantly, MP measurements were conducted at nanomolar protein concentrations, below the threshold for macroscopic condensate formation. Under high AmAc conditions, tau was predominantly monomeric, consistent with its fully dispersed state at high salt (Figure 2d). At intermediate AmAc concentrations, MP revealed the presence of tau dimers and trimers. At the lowest AmAc concentration, tau appeared mainly monomeric, potentially due to the formation of polydisperse clusters that precede macroscopic condensates ^33^. To further probe the interactions that stabilize tau assemblies, we perturbed pre-formed droplets with 1,6-hexanediol or formic acid (Figure S2c). Only formic acid dissolved the droplets, showing that electrostatic interactions, rather than hydrophobic contacts, dominate tau LLPS under low-salt conditions.

The combined insights from microscopy, native MS, IMMS, and MP align with the prevalent model for salt-controlled tau LLPS, in which decreasing ionic strength reorganizes tau’s conformational ensemble through electrostatically driven compaction, assembly formation, and ultimately LLPS (Figure S2d) ^34^. The approach thus enables us to directly observe how interactions with potential binding partners are affected by the LLPS regime.

### BRICHOS binds the proline-rich region of tau selectively under LLPS conditions

To assess how LLPS-dependent conformational changes influence protein binding to tau, we first analyzed tau–BRICHOS interactions. Native MS of 20 μM tau and 10 μM BRICHOS in the presence of 100 mM AmAc, *i*.*e*. non-LLPS conditions, show no interactions between both proteins (Figure 3a). However, when the AmAc concentration was reduced to 2.5 mM, *i*.*e*. promoting LLPS conditions, we detected prominent peaks corresponding in mass to a 1:1 complex between tau and BRICHOS. Importantly, no other stoichiometries were observed (Figure 3a). To determine whether the dependency of the interaction on salt concentration is detectable in solution, we turned to MP. Although the free BRICHOS protein (14 kDa) is below the MP detection threshold, we observe a broadening of the tau monomer peak (45.7 kDa) in the presence of BRICHOS in MilliQ, indicating complex formation (Figure 3b). No peak broadening was observed when the MP measurements were performed in 100 mM AmAc.

**Figure 3.**
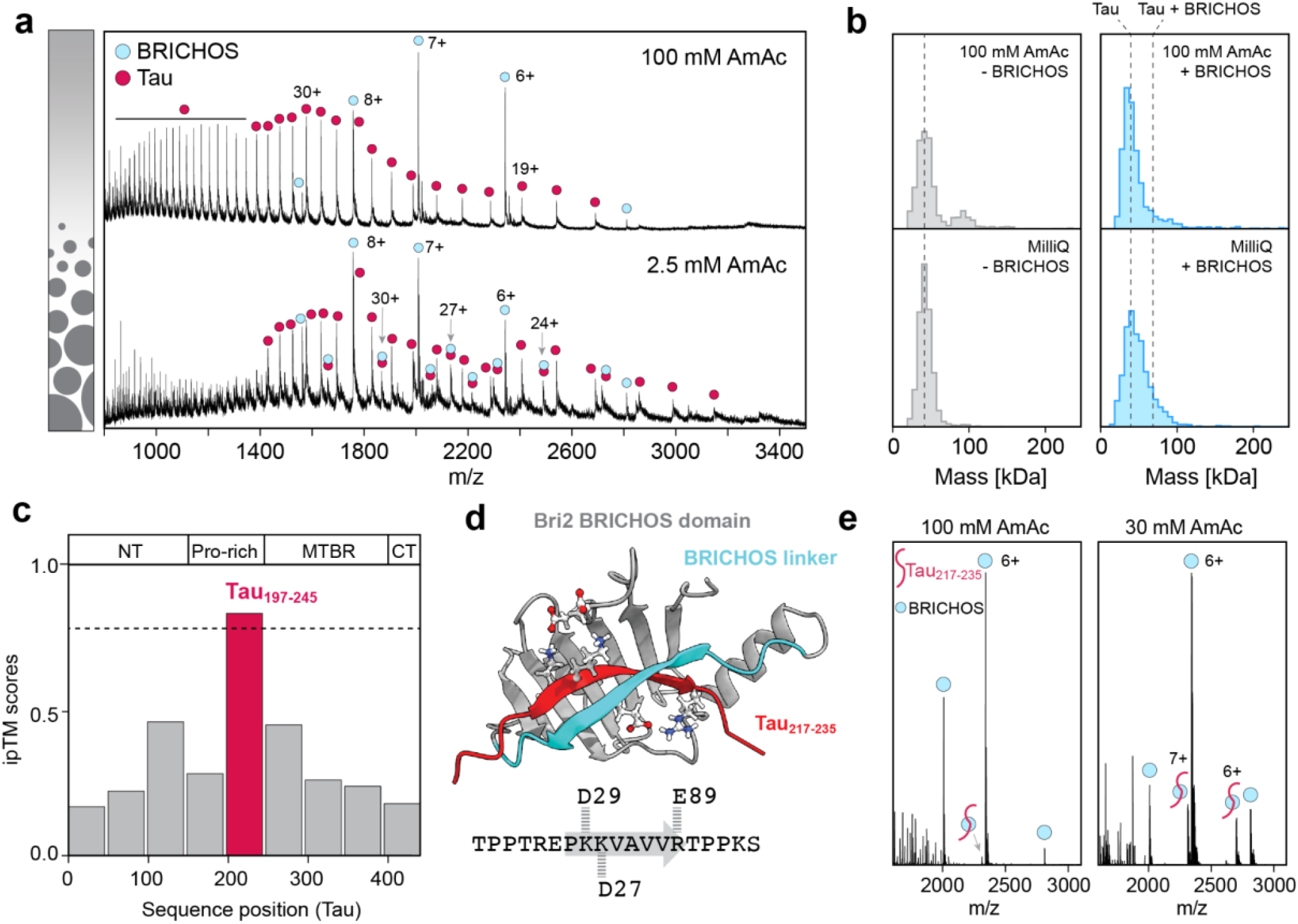
Tau-BRICHOS interaction and binding site determination under LLPS-promoting conditions. (a) Native mass spectra of 20 μM tau (red) with 10 μM BRICHOS (blue) in 100 mM and 2.5 mM AmAc pH 8. BRICHOS binds to tau only under low salt concentration. (b) MP histograms of 20 nM tau with 200 nM BRICHOS in high and low AmAc concentrations. (c) ipTM scores for BRICHOS binding to 49 residue peptides of tau, predicted with AlphaFold2. For peptide 5 (red; tau_197–245_) a high confidence interaction was predicted (ipTM > 0.8). (d) Structure of BRICHOS (gray with blue linker) in complex with tau_217-235_ (red). The interacting charged residues are shown as stick representation. The tau sequence is shown below with the BRICHOS-interacting residues indicated in gray. (e) Native mass spectra of 25 μM tau_217-235_ (red) with 5 μM BRICHOS (blue) in 100 mM and 30 mM AmAc pH 8 show BRICHOS binding to tau_217-235_ only at low salt concentration.

These findings raise the question of how changes in salt concentration that drive tau LLPS concomitantly control the association of tau and BRICHOS. As the first step, we mapped the tau region responsible for BRICHOS binding. We divided the tau sequence into nine equal-length peptides (49 residues) and predicted their interactions with BRICHOS using AlphaFold2 ^35^. AlphaFold2 was chosen as AF2-based approaches have been shown to be effective in predicting weak and transient protein–protein interactions, making it well suited for mapping candidate BRICHOS-binding regions within tau ^36^. Only peptide 5 (residues 197–245), located in the proline-rich region, resulted in an ipTM score above 0.8, indicating high-confidence binding (Figure 3c). Predicting the BRICHOS structure with this peptide markedly increased the pLDDT score of the otherwise disordered BRICHOS linker (Figure S3a). The resulting structure suggests a complementary β-strand formation between the tau peptide and the BRICHOS linker region (Figure 3d), a binding mode previously reported for other BRICHOS–client interactions ^37^.

Next, we sought to experimentally validate the predicted binding site by performing native MS of BRICHOS with a 19-residue peptide (residues 217–235) covering the predicted interacting region of tau. At high AmAc concentrations, no interaction was detected, whereas a 1:1 BRICHOS–tau_217–235_ complex could be observed at low AmAc concentrations (Figure 3e), reflecting the observations for full-length tau. To identify potential causes for the salt dependence, we conducted molecular dynamics (MD) simulations of the tau_217–235_ peptide. Simulations with 200 mM AmAc produced preferentially compact conformations. Reducing the AmAc concentration promoted a more extended geometry (Figure S3b). The conformational shift likely stems from reduced charge screening in the absence of salt, increasing electrostatic repulsion between positively charged residues and positioning them to form favorable salt bridges with acidic residues in BRICHOS, thereby promoting binding under LLPS-promoting conditions. In summary, our data suggest that BRICHOS binds to tau under LLPS-promoting conditions *via* a salt-dependent conformational relay encompassing residues 223–231 in the proline-rich region.

To find out whether the BRICHOS-tau interaction is specific for BRICHOS from the Bri2 protein, we also tested the BRICHOS domain from the lung surfactant proprotein C (pro-SPC). proSP-C BRICHOS recognizes hydrophobic sequences with β-strand propensity and is a potent inhibitor of amyloid β aggregation *in vitro* ^38,39,37^. Comparison of the experimental proSP-C BRICHOS structure with the predicted Bri2 BRICHOS structure confirms an essentially identical fold (Figure S3c) ^40^. However, AlphaFold2-docking of the tau peptides into proSP-C BRICHOS does not yield models with increased ipTM scores for the tau fragments (Figure S3d). Native MS analysis showed no pronounced binding of tau_217–235_ to proSP-C BRICHOS regardless of AmAc concentration (Figure S3e). We conclude that the LLPS-associated BRICHOS-tau interaction is specific for the BRICHOS domain from the Bri2 protein.

### BRICHOS interferes with tubulin binding to tau under LLPS conditions

Considering the possibility that BRICHOS recruitment is controlled by LLPS-dependent conformational changes in tau, we then asked whether these interactions could provide a mechanistic explanation into the BRICHOS-dependent inhibition of microtubuli assembly in tau condensates (Figure 1). In the high-resolution structure of a microtubule and a tau fragment, the microtubule-binding repeats (MTBR) of tau tightly associate with the tubulin surface, whereas the N-terminally adjacent proline-rich region (PRR) encompassing the putative BRICHOS binding site is partially detached (Figure 4a) ^41^. Expanding on the AlphaFold predictions of tau and BRICHOS (Figure 3c, d), we predicted pairwise complexes between tau residues 200–275 (spanning the PRR and the first MTBR) and BRICHOS or tubulin, and monitored the resulting pLDDT scores using AlphaFold3 (Figure 4a). Since the tau segment in isolation is disordered, an increase in pLDDT score suggests docking into the folded binding partner. For BRICHOS, the predictions suggest binding to the PRR, consistent with our experimental data, and tubulin was suggested to bind to the MTBR, in agreement with the high-resolution structure (Figure 4a, Figure S4a). However, when we predicted a ternary complex composed of tau, BRICHOS, and tubulin, only the tubulin-binding region of tau showed increased pLDDT scores, suggesting that BRICHOS and tubulin interactions at their adjacent binding sites are mutually exclusive.

**Figure 4.**
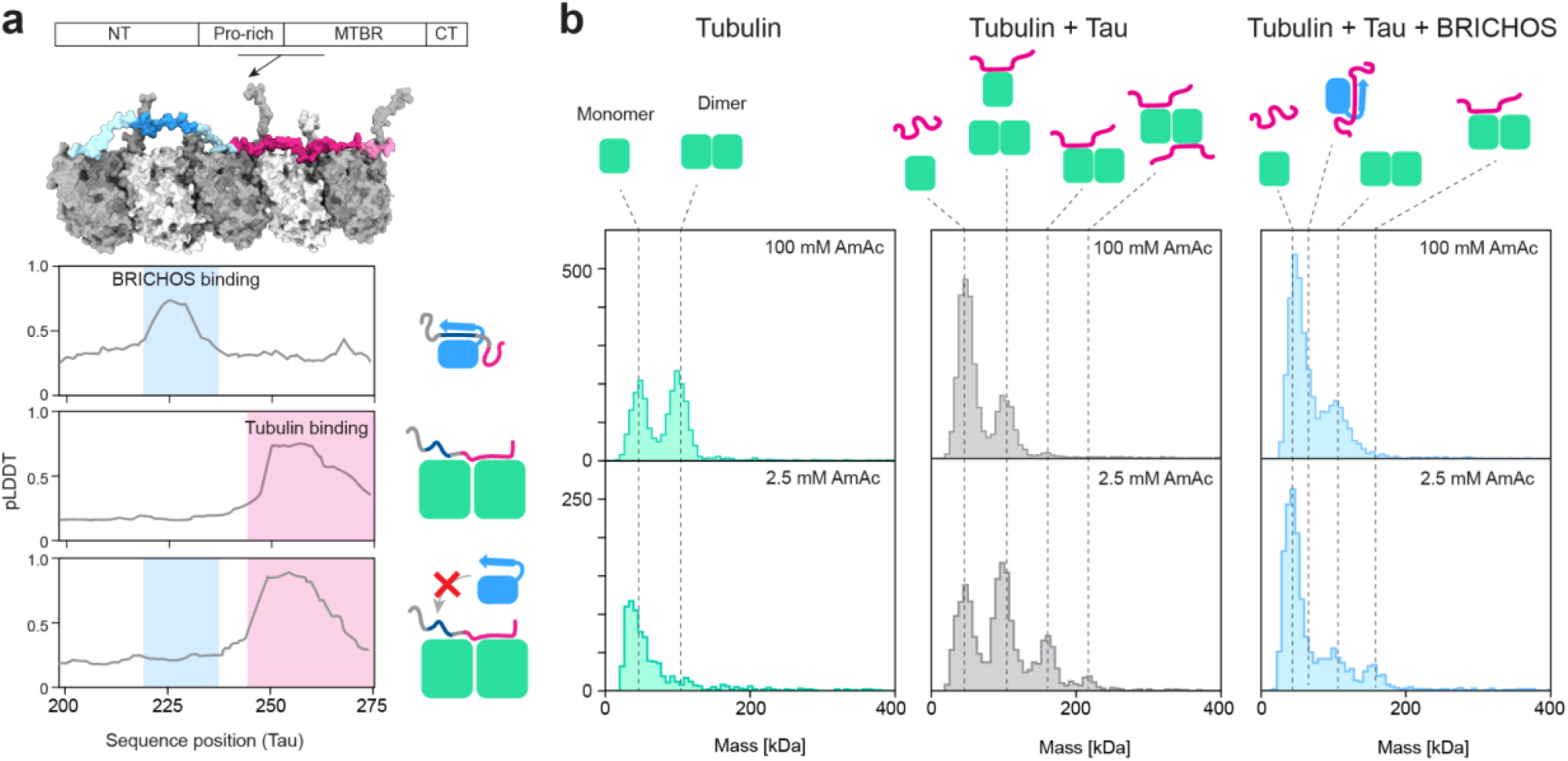
Competitive binding of BRICHOS and tubulin to tau under LLPS-promoting conditions. (a) Structure of a tubulin oligomer in complex with tau residues 202-281 (PDB ID 7PQC) ^41^. Tubulin is shown in grey, the BRICHOS and tubulin binding regions of tau are highlighted in blue and pink, respectively. Below: AlphaFold3 pLDDT scores for BRICHOS and tubulin complexes with the relevant region of tau (residues 197-274) suggest closely adjacent binding sites. Tau and tubulin binding locally increase the pLDDT score of the respective tau region. In the ternary complex prediction (bottom), only the tubulin binding region shows increased scores, suggesting that tubulin and BRICHOS interactions are mutually exclusive. (b) MP histograms of tau (20 nM) and tubulin (5 nM) with and without BRICHOS (200 nM) under LLPS and non-LLPS conditions. Tubulin monomers and dimers are detected at 100 mM AmAc and only monomers at 2.5 mM AmAc. Tubulin with tau leads to complex formation seen as higher-order oligomers. BRICHOS interferes with tau-tubulin interactions, as evident from the lower oligomer count.

To test this hypothesis experimentally, we performed MP on tau and tubulin in the presence and absence of BRICHOS (Figure 4b). Tau (46 kDa) and tubulin (55 kDa) have similar molecular weights, making individual peaks difficult to discern. However, tubulin alone formed monomers and dimers at high AmAc concentrations, and predominantly monomers under LLPS-promoting conditions (Figure 4b, left column). Upon addition of tau, higher oligomeric species appeared specifically under LLPS conditions that were not observed for either protein alone. These new species have a molecular weight of approximately 160 and 210 kDa, consistent with a tubulin dimer binding one or two tau molecules (Figure 4b, middle column). Co-incubation of tau-tubulin complexes with excess BRICHOS largely abolished these higher-order species, yielding primarily monomeric tau and tubulin species (Figure 4b, right column). No interactions between tubulin and BRICHOS could be detected (Figure S4b). Taken together, the AlphaFold3 predictions and MP data lead us to conclude that BRICHOS and tubulin compete for binding to tau under LLPS-promoting conditions. This competition mechanism is consistent with the reduced ability of tau condensates to promote microtubule assembly in the presence of BRICHOS (Figure 1b), strongly suggesting that the interactions observed by MS and MP can take place in the dense phase.

## Discussion

Understanding how interactions in IDPs are regulated remains a major challenge, since they populate dynamic ensembles and engage partners through transient, multivalent interactions. LLPS adds another layer of complexity by reorganizing these ensembles into locally concentrated, dynamically exchanging populations with potentially altered conformational landscapes. In this work, we establish an integrated framework combining MS, MP, computational modeling and microscopy that captures LLPS-dependent conformational changes of tau and how it affects client engagement in multicomponent systems.

Using native IMMS and mass photometry, we observe that LLPS-promoting conditions reshape tau’s conformational landscape: the ensemble shifts toward more compact monomeric states driven by electrostatic interactions, and transient oligomers appear at intermediate ionic strengths. Similar changes have been identified in other IDPs whose phase separation is driven by a balance between sequence-encoded propensities for collapse and multivalent intermolecular contacts ^42^. This concept raises the possibility that salt-driven tau LLPS not only assembles the protein, but also modulates the accessibility of weak binding interfaces which we can resolve with our approach.

Here, we show that restructuring the conformational ensemble of tau through LLPS-promoting conditions has direct consequences for its interactions with protein clients. NMR studies have demonstrated that tau has a propensity to adopt local secondary structures, including transient β-structure-like segments and regions with α-helical character ^11^. Interestingly, the region within the proline-rich domain that we identify as a BRICHOS-binding site is among those capable of sampling β-extended conformations. In addition to the conformational plasticity of tau, a previous study indicates that flexible regions within the Bri2 BRICHOS domain can sample alternative conformations, providing adaptable interaction surfaces for binding structurally diverse clients ^43^. Such flexibility on both sides may stabilize transient interactions under LLPS-promoting conditions, positioning tau to engage BRICHOS through β-strand complementation. Importantly, this binding-competent state is captured experimentally through native MS, mass photometry, and computational modeling. Notably, BRICHOS–tau interactions have previously been observed under non-LLPS conditions, indicating that phase separation is not strictly required for binding ^21^. However, LLPS reorganizes tau’s conformational ensemble and promotes co-localization of tau and BRICHOS within the dense phase, such that condensate formation and chaperone engagement converge. Given the conformational heterogeneity of tau and the flexible nature of BRICHOS, there are likely additional low-affinity interaction sites that contribute to their multivalent interaction landscape.

Previous studies showed that the conformational plasticity of tau can give rise to seeds with distinct aggregation characteristics associated with different tauopathies ^44^. These conformations expose specific amyloidogenic motifs in the MTBR and the proline-rich region, which can be recognized by chaperones that stabilize monomeric tau and reduce aggregation propensity. A prominent example is Clusterin, which forms a membrane-associated complex that recognizes the MTBR of tau *via* its disordered tails ^45^. J-domain proteins bind aggregation-prone tau monomers and recruit HSP70 for refolding. Some members of this family, like DnaJB1 and DnaJB2, additionally recognize tau oligomers and help in their disaggregation ^46^. The Ca^2+^ activated chaperone S100B specifically acts on tau condensates, shifting the equilibrium towards the soluble form and thus reducing the risk of aggregation from tau condensates ^47^. Our finding that chaperone binding to tau is enhanced under LLPS-promoting conditions provides a model for how such interactions may be regulated. The fact that BRICHOS-tau interactions directly impact tubulin assembly leads us to propose a relay mechanism in which phase separation modulates the balance between tubulin-engaged and chaperone-engaged tau populations. Interestingly, the BRICHOS-binding segment covers multiple disease-associated phosphorylation sites in tau, including Threonine 231 and Serine 235. Phosphorylation at these sites controls the ability of tau to mediate tubulin assembly ^48^, indicating a regulatory circuit that overlaps with BRICHOS interactions.

A better understanding how chaperones recognize LLPS-dependent conformations, and how these interactions impact condensate function, opens new opportunities for the development of therapeutics. Considering that tau aggregation can be initiated by LLPS ^49^, the ability to identify and target specific functional states offers a strategic advantage. The experimental framework presented here has the potential to reveal how disease-associated post-translational modifications, ligands, or mutations alter the LLPS-dependent interaction landscape of tau, and identify binding interfaces that represent vulnerable nodes which can be targeted therapeutically. More broadly, our approach offers a path to dissect how LLPS can reshape conformational landscapes and interaction specificities across diverse IDPs.

## Acknowledgements

ML is supported by a KI faculty-funded Career Position, a Cancerfonden Project grant, a VR Research Environment Grant, a Consolidator Grant from the Swedish Society for Medical Research (SSMF) and a Project Grant from the Knut och Alice Wallenberg Foundation (KAW). AA is supported by a Consolidator Grant from Karolinska Institutet, the Swedish research council, the Swedish Alzheimer Foundation, the Swedish Brain Foundation and CIMED. CM is supported by the Swedish Brain Foundation, Petrus and Augusta Hedlund Foundation, Astrid and David Hageléns Foundation, and the Åhléns Foundation.

## Notes

### Competing Interest Statement

The authors have declared no competing interest.

### Summary of Updates

Author list and affiliations were updated; minor phrasing changes

## References

(1) Holehouse, A. S.; Kragelund, B. B. The Molecular Basis for Cellular Function of Intrinsically Disordered Protein Regions. Nat. Rev. Mol. Cell Biol. 2024, 25 (3), 187–211. 10.1038/s41580-023-00673-0.

(2) Keppel, T. R.; Howard, B. A.; Weis, D. D. Mapping Unstructured Regions and Synergistic Folding in Intrinsically Disordered Proteins with Amide H/D Exchange Mass Spectrometry. Biochemistry 2011, 50 (40), 8722–8732. 10.1021/bi200875p.

(3) Wang, J.; Choi, J.-M.; Holehouse, A. S.; Lee, H. O.; Zhang, X.; Jahnel, M.; Maharana, S.; Lemaitre, R.; Pozniakovsky, A.; Drechsel, D.; Poser, I.; Pappu, R. V.; Alberti, S.; Hyman, A. A. A Molecular Grammar Governing the Driving Forces for Phase Separation of Prion-like RNA Binding Proteins. Cell 2018, 174 (3), 688-699.e16. 10.1016/j.cell.2018.06.006.

(4) Rekhi, S.; Garcia, C. G.; Barai, M.; Rizuan, A.; Schuster, B. S.; Kiick, K. L.; Mittal, J. Expanding the Molecular Language of Protein Liquid–Liquid Phase Separation. Nat. Chem. 2024, 16 (7), 1113–1124. 10.1038/s41557-024-01489-x.

(5) Yamazaki, H.; Takagi, M.; Kosako, H.; Hirano, T.; Yoshimura, S. H. Cell Cycle-Specific Phase Separation Regulated by Protein Charge Blockiness. Nat. Cell Biol. 2022, 24 (5), 625–632. 10.1038/s41556-022-00903-1.

(6) Martin, E. W.; Holehouse, A. S.; Peran, I.; Farag, M.; Incicco, J. J.; Bremer, A.; Grace, C. R.; Soranno, A.; Pappu, R. V.; Mittag, T. Valence and Patterning of Aromatic Residues Determine the Phase Behavior of Prion-like Domains. Science 2020, 367 (6478), 694–699. 10.1126/science.aaw8653.

(7) Galvanetto, N.; Ivanović, M. T.; Del Grosso, S. A.; Chowdhury, A.; Sottini, A.; Nettels, D.; Best, R. B.; Schuler, B. Material Properties of Biomolecular Condensates Emerge from Nanoscale Dynamics. Proc. Natl. Acad. Sci. 2025, 122 (23), e2424135122. 10.1073/pnas.2424135122.

(8) Banani, S. F.; Lee, H. O.; Hyman, A. A.; Rosen, M. K. Biomolecular Condensates: Organizers of Cellular Biochemistry. Nat. Rev. Mol. Cell Biol. 2017, 18 (5), 285–298. 10.1038/nrm.2017.7.

(9) Nott, T. J.; Petsalaki, E.; Farber, P.; Jervis, D.; Fussner, E.; Plochowietz, A.; Craggs, T. D.; Bazett-Jones, D. P.; Pawson, T.; Forman-Kay, J. D.; Baldwin, A. J. Phase Transition of a Disordered Nuage Protein Generates Environmentally Responsive Membraneless Organelles. Mol. Cell 2015, 57 (5), 936–947. 10.1016/j.molcel.2015.01.013.

(10) Wegmann, S.; Eftekharzadeh, B.; Tepper, K.; Zoltowska, K. M.; Bennett, R. E.; Dujardin, S.; Laskowski, P. R.; MacKenzie, D.; Kamath, T.; Commins, C.; Vanderburg, C.; Roe, A. D.; Fan, Z.; Molliex, A. M.; Hernandez-Vega, A.; Muller, D.; Hyman, A. A.; Mandelkow, E.; Taylor, J. P.; Hyman, B. T. Tau Protein Liquid–Liquid Phase Separation Can Initiate Tau Aggregation. EMBO J. 2018, 37 (7), EMBJ201798049. 10.15252/embj.201798049.

(11) Mukrasch, M. D.; Bibow, S.; Korukottu, J.; Jeganathan, S.; Biernat, J.; Griesinger, C.; Mandelkow, E.; Zweckstetter, M. Structural Polymorphism of 441-Residue Tau at Single Residue Resolution. PLOS Biol. 2009, 7 (2), e1000034. 10.1371/journal.pbio.1000034.

(12) Boyko, S.; Qi, X.; Chen, T.-H.; Surewicz, K.; Surewicz, W. K. Liquid-Liquid Phase Separation of Tau Protein: The Crucial Role of Electrostatic Interactions. J. Biol. Chem. 2019, 294 (29), 11054–11059. 10.1074/jbc.AC119.009198.

(13) Hernández-Vega, A.; Braun, M.; Scharrel, L.; Jahnel, M.; Wegmann, S.; Hyman, B. T.; Alberti, S.; Diez, S.; Hyman, A. A. Local Nucleation of Microtubule Bundles through Tubulin Concentration into a Condensed Tau Phase. Cell Rep. 2017, 20 (10), 2304–2312. 10.1016/j.celrep.2017.08.042.

(14) Kadavath, H.; Jaremko, M.; Jaremko, Ł.; Biernat, J.; Mandelkow, E.; Zweckstetter, M. Folding of the Tau Protein on Microtubules. Angew. Chem. Int. Ed. 2015, 54 (35), 10347–10351. 10.1002/anie.201501714.

(15) Cario, A.; Berger, C. L. Tau, Microtubule Dynamics, and Axonal Transport: New Paradigms for Neurodegenerative Disease. BioEssays 2023, 45 (8), 2200138. 10.1002/bies.202200138.

(16) Fitzpatrick, A. W. P.; Falcon, B.; He, S.; Murzin, A. G.; Murshudov, G.; Garringer, H. J.; Crowther, R. A.; Ghetti, B.; Goedert, M.; Scheres, S. H. W. Cryo-EM Structures of Tau Filaments from Alzheimer’s Disease. Nature 2017, 547 (7662), 185–190. 10.1038/nature23002.

(17) Goedert, M.; Spillantini, M. G. Propagation of Tau Aggregates. Mol. Brain 2017, 10 (1), 18. 10.1186/s13041-017-0298-7.

(18) Kanaan, N. M.; Hamel, C.; Grabinski, T.; Combs, B. Liquid-Liquid Phase Separation Induces Pathogenic Tau Conformations in Vitro. Nat. Commun. 2020, 11 (1), 2809. 10.1038/s41467-020-16580-3.

(19) Alberti, S.; Gladfelter, A.; Mittag, T. Considerations and Challenges in Studying Liquid-Liquid Phase Separation and Biomolecular Condensates. Cell 2019, 176 (3), 419–434. 10.1016/j.cell.2018.12.035.

(20) Leder, A.; Mas, G.; Szentgyörgyi, V.; Jakob, R. P.; Maier, T.; Spang, A.; Hiller, S. A Multichaperone Condensate Enhances Protein Folding in the Endoplasmic Reticulum. Nat. Cell Biol. 2025, 27 (9), 1422–1430. 10.1038/s41556-025-01730-w.

(21) Mörman, C.; Leppert, A.; Pizzirusso, G.; Zheng, Z.; Sun, X.; Kumar, R.; Biverstål, H.; Landreh, M.; Johansson, J.; Arroyo-Garcia, L. E.; Luo, J.; Chen, G.; Abelein, A. Chaperone-Mediated Regulation of Tau Phase Separation, Fibrillation, and Toxicity. J. Am. Chem. Soc. 2025, 147 (27), 23504–23518. 10.1021/jacs.5c01369.

(22) Seidler, P. M.; Murray, K. A.; Boyer, D. R.; Ge, P.; Sawaya, M. R.; Hu, C. J.; Cheng, X.; Abskharon, R.; Pan, H.; DeTure, M. A.; Williams, C. K.; Dickson, D. W.; Vinters, H. V.; Eisenberg, D. S. Structure-Based Discovery of Small Molecules That Disaggregate Alzheimer’s Disease Tissue Derived Tau Fibrils in Vitro. Nat. Commun. 2022, 13 (1), 5451. 10.1038/s41467-022-32951-4.

(23) Powell, W. C.; Nahum, M.; Pankratz, K.; Herlory, M.; Greenwood, J.; Poliyenko, D.; Holland, P.; Jing, R.; Biggerstaff, L.; Stowell, M. H. B.; Walczak, M. A. Post-Translational Modifications Control Phase Transitions of Tau. ACS Cent. Sci. 2024, 10 (11), 2145–2161. 10.1021/acscentsci.4c01319.

(24) Shimozawa, M.; Tigro, H.; Biverstål, H.; Shevchenko, G.; Bergquist, J.; Moaddel, R.; Johansson, J.; Nilsson, P. Identification of Cytoskeletal Proteins as Binding Partners of Bri2 BRICHOS Domain. Mol. Cell. Neurosci. 2023, 125, 103843. 10.1016/j.mcn.2023.103843.

(25) Wohlschlegel, J.; Argentini, M.; Michiels, C.; Letellier, C.; Forster, V.; Condroyer, C.; He, Z.; Thuret, G.; Zeitz, C.; Léger, T.; Audo, I. First Identification of ITM2B Interactome in the Human Retina. Sci. Rep. 2021, 11 (1), 17210. 10.1038/s41598-021-96571-6.

(26) Chen, G.; Andrade-Talavera, Y.; Tambaro, S.; Leppert, A.; Nilsson, H. E.; Zhong, X.; Landreh, M.; Nilsson, P.; Hebert, H.; Biverstål, H.; Fisahn, A.; Abelein, A.; Johansson, J. Augmentation of Bri2 Molecular Chaperone Activity against Amyloid-β Reduces Neurotoxicity in Mouse Hippocampus in Vitro. Commun. Biol. 2020, 3 (1), 32. 10.1038/s42003-020-0757-z.

(27) Sahin, C.; Leppert, A.; Landreh, M. Advances in Mass Spectrometry to Unravel the Structure and Function of Protein Condensates. Nat. Protoc. 2023, 18 (12), 3653–3661. 10.1038/s41596-023-00900-0.

(28) Sahin, C.; Motso, A.; Gu, X.; Feyrer, H.; Lama, D.; Arndt, T.; Rising, A.; Gese, G. V.; Hällberg, B. M.; Marklund, Erik. G.; Schafer, N. P.; Petzold, K.; Teilum, K.; Wolynes, P. G.; Landreh, M. Mass Spectrometry of RNA-Binding Proteins during Liquid–Liquid Phase Separation Reveals Distinct Assembly Mechanisms and Droplet Architectures. J. Am. Chem. Soc. 2023, 145 (19), 10659–10668. 10.1021/jacs.3c00932.

(29) Li, J.; Santambrogio, C.; Brocca, S.; Rossetti, G.; Carloni, P.; Grandori, R. Conformational Effects in Protein Electrospray-ionization Mass Spectrometry. Mass Spectrom. Rev. 2016, 35 (1), 111–122. 10.1002/mas.21465.

(30) Osterholz, H.; Stevens, A.; Abramsson, M. L.; Lama, D.; Brackmann, K.; Rising, A.; Elofsson, A.; Marklund, E. G.; Deindl, S.; Leppert, A.; Landreh, M. Native Mass Spectrometry Captures the Conformational Plasticity of Proteins with Low-Complexity Domains. JACS Au 2025, 5 (1), 281–290. 10.1021/jacsau.4c00961.

(31) Beveridge, R.; Migas, L. G.; Das, R. K.; Pappu, R. V.; Kriwacki, R. W.; Barran, P. E. Ion Mobility Mass Spectrometry Uncovers the Impact of the Patterning of Oppositely Charged Residues on the Conformational Distributions of Intrinsically Disordered Proteins. J. Am. Chem. Soc. 2019, 141 (12), 4908–4918. 10.1021/jacs.8b13483.

(32) Konermann, L. Addressing a Common Misconception: Ammonium Acetate as Neutral pH “Buffer” for Native Electrospray Mass Spectrometry. J. Am. Soc. Mass Spectrom. 2017, 28 (9), 1827–1835. 10.1007/s13361-017-1739-3.

(33) Ray, S.; Mason, T. O.; Boyens-Thiele, L.; Farzadfard, A.; Larsen, J. A.; Norrild, R. K.; Jahnke, N.; Buell, A. K. Mass Photometric Detection and Quantification of Nanoscale α-Synuclein Phase Separation. Nat. Chem. 2023, 15 (9), 1306–1316. 10.1038/s41557-023-01244-8.

(34) Wen, J.; Tang, Y.; Sneideris, T.; Ausserwöger, H.; Hong, L.; Knowles, T. P. J.; Perrett, S.; Wei, G.; Wu, S. Direct Observation of the Conformational Transitions in Tau and Their Correlation with Phase Behavior. JACS Au 2025, 5 (9), 4268–4280. 10.1021/jacsau.5c00625.

(35) Mirdita, M.; Schütze, K.; Moriwaki, Y.; Heo, L.; Ovchinnikov, S.; Steinegger, M. ColabFold: Making Protein Folding Accessible to All. Nat. Methods 2022, 19 (6), 679–682. 10.1038/s41592-022-01488-1.

(36) Chojnowski, G. gapTrick—Structural Characterization of Protein–Protein Interactions Using AlphaFold. Bioinformatics 2025, 41 (9), btaf532. 10.1093/bioinformatics/btaf532.

(37) Leppert, A.; Poska, H.; Landreh, M.; Abelein, A.; Chen, G.; Johansson, J. A New Kid in the Folding Funnel: Molecular Chaperone Activities of the BRICHOS Domain. Protein Sci. 2023, 32 (6), e4645. 10.1002/pro.4645.

(38) Nerelius, C.; Gustafsson, M.; Nordling, K.; Larsson, A.; Johansson, J. Anti-Amyloid Activity of the C-Terminal Domain of proSP-C against Amyloid β-Peptide and Medin. Biochemistry 2009, 48 (17), 3778–3786. 10.1021/bi900135c.

(39) Johansson, H.; Nerelius, C.; Nordling, K.; Johansson, J. Preventing Amyloid Formation by Catching Unfolded Transmembrane Segments. J. Mol. Biol. 2009, 389 (2), 227–229. 10.1016/j.jmb.2009.04.021.

(40) Willander, H.; Askarieh, G.; Landreh, M.; Westermark, P.; Nordling, K.; Keränen, H.; Hermansson, E.; Hamvas, A.; Nogee, L. M.; Bergman, T.; Saenz, A.; Casals, C.; Åqvist, J.; Jörnvall, H.; Berglund, H.; Presto, J.; Knight, S. D.; Johansson, J. High-Resolution Structure of a BRICHOS Domain and Its Implications for Anti-Amyloid Chaperone Activity on Lung Surfactant Protein C. Proc. Natl. Acad. Sci. 2012, 109 (7), 2325–2329. 10.1073/pnas.1114740109.

(41) Brotzakis, Z. F.; Lindstedt, P. R.; Taylor, R. J.; Rinauro, D. J.; Gallagher, N. C. T.; Bernardes, G. J. L.; Vendruscolo, M. A Structural Ensemble of a Tau-Microtubule Complex Reveals Regulatory Tau Phosphorylation and Acetylation Mechanisms. ACS Cent. Sci. 2021, 7 (12), 1986–1995. 10.1021/acscentsci.1c00585.

(42) Krainer, G.; Welsh, T. J.; Joseph, J. A.; Espinosa, J. R.; Wittmann, S.; de Csilléry, E.; Sridhar, A.; Toprakcioglu, Z.; Gudiškytė, G.; Czekalska, M. A.; Arter, W. E.; Guillén-Boixet, J.; Franzmann, T. M.; Qamar, S.; George-Hyslop, P. S.; Hyman, A. A.; Collepardo-Guevara, R.; Alberti, S.; Knowles, T. P. J. Reentrant Liquid Condensate Phase of Proteins Is Stabilized by Hydrophobic and Non-Ionic Interactions. Nat. Commun. 2021, 12 (1), 1085. 10.1038/s41467-021-21181-9.

(43) Kumar, R.; Le Marchand, T.; Adam, L.; Bobrovs, R.; Chen, G.; Fridmanis, J.; Kronqvist, N.; Biverstål, H.; Jaudzems, K.; Johansson, J.; Pintacuda, G.; Abelein, A. Identification of Potential Aggregation Hotspots on Aβ42 Fibrils Blocked by the Anti-Amyloid Chaperone-like BRICHOS Domain. Nat. Commun. 2024, 15 (1), 965. 10.1038/s41467-024-45192-4.

(44) Hou, Z.; Chen, D.; Ryder, B. D.; Joachimiak, L. A. Biophysical Properties of a Tau Seed. Sci. Rep. 2021, 11 (1), 13602. 10.1038/s41598-021-93093-z.

(45) Yuste-Checa, P.; Carvajal, A. I.; Mi, C.; Paatz, S.; Hartl, F. U.; Bracher, A. Structural Analyses Define the Molecular Basis of Clusterin Chaperone Function. Nat. Struct. Mol. Biol. 2025, 32 (10), 2035–2045. 10.1038/s41594-025-01631-4.

(46) Ryder, B. D.; Wydorski, P. M.; Hou, Z.; Joachimiak, L. A. Chaperoning Shape-Shifting Tau in Disease. Trends Biochem. Sci. 2022, 47 (4), 301–313. 10.1016/j.tibs.2021.12.009.

(47) Moreira, G. G.; Gomes, C. M. Tau Liquid–Liquid Phase Separation Is Modulated by the Ca^2+^-switched Chaperone Activity of the S100B Protein. J. Neurochem. 2023, 166 (1), 76–86. 10.1111/jnc.15756.

(48) Savastano, A.; Flores, D.; Kadavath, H.; Biernat, J.; Mandelkow, E.; Zweckstetter, M. Disease-Associated Tau Phosphorylation Hinders Tubulin Assembly within Tau Condensates. Angew. Chem. Int. Ed. 2021, 60 (2), 726–730. 10.1002/anie.202011157.

(49) Kanaan, N. M.; Hamel, C.; Grabinski, T.; Combs, B. Liquid-Liquid Phase Separation Induces Pathogenic Tau Conformations in Vitro. Nat. Commun. 2020, 11 (1), 2809. 10.1038/s41467-020-16580-3.

