## Supplementary for "An electrostatic relay connects client binding and liquid-liquid phase separation of tau"

**Methods**

**Protein expression and purification**

Tau protein was expressed and purified as described previously, with small alterations [1]. Shortly, the protein was expressed in BL21 (DE3) *E. coli* cells for 2.5 h with 0.4 mM IPTG. The harvested cells were resuspended in 50 mM $NaP_i$, 1 mM EDTA pH6.4 with 1x protease inhibitor cocktail (cOmplete,EDTA-free, Roche) and disrupted by sonication (6 min, 2s on/8s off, 40%). Cell debris was removed by centrifugation and the supernatant was heat treated and centrifuged again to remove the insoluble fraction. After cation exchange, size exclusion (Superdex 200 PG 26/600) was used for further purification and buffer exchange to 20 mM NaPi, 0.2 mM EDTA, pH 7.4.

The monomeric variant of the recombinant human Bri2 BRICHOS domain carrying an arginine-to-glutamate substitution at position 221 (R221E) was produced as described previously [2]. Briefly, the protein was expressed recombinantly in *Escherichia coli* BL21 (DE3) cells as a His-NT*-tag fusion by induction with 0.5 mM IPTG at an $OD_{600}$ of 0.7–0.9 and incubated overnight at 20 °C. Cells were harvested by centrifugation, resuspended in 20 mM Tris-HCl, pH 8.0 supplemented with EDTA-free protease inhibitor cocktail (cOmplete, Roche), and lysed by sonication. The protein was purified by Ni–NTA affinity chromatography, followed by size exclusion chromatography using a Superdex 200 column (GE Healthcare). The monomeric fusion protein fraction was collected, the His-tag was removed by thrombin cleavage, and the sample was subjected to a second Ni–NTA step to remove the released tag and uncleaved protein. Monomeric Bri2 BRICHOS R221E was subsequently isolated by size exclusion chromatography using a Superdex 75 column (GE Healthcare) in 20 mM NaPi, 0.2 mM EDTA, pH 8, aliquoted, and stored at −20 °C until further use.

The monomeric variant of the recombinant human proSP-C BRICHOS domain (residues 59–197 of human proSP-C; UniProt KB accession number P11686) carrying a threonine-to-arginine substitution at position 187 (T187R) was produced as described previously [3]. Briefly, the protein was expressed recombinantly in *Escherichia coli* Origami (DE3) pLysS cells as a thioredoxin–His-tag fusion by induction with 0.5 mM IPTG at an $OD_{600}$ of 0.7–0.9 and incubated for 6 h at 37 °C. Cells were harvested by centrifugation, resuspended in 20 mM Tris-HCl, pH 8.0 supplemented with EDTA-free protease inhibitor cocktail (cOmplete, Roche), and lysed by sonication. The protein was purified under denaturing conditions (2 M Urea in 20 mM Tris, pH 8) by Ni–NTA affinity chromatography. Following refolding by dialysis against 20 mM Tris, pH 8, the fusion tag was removed by thrombin cleavage, and uncleaved protein and tag fragments were removed by a second Ni–NTA step. The monomeric fraction was purified by size exclusion chromatography using a Superdex 75 PG 26/600 column (GE Healthcare) in 20 mM NaPi, 0.2 mM EDTA, pH 8. Protein concentrations were determined by absorbance at 280 nm.

**Protein labeling**

Prior to fluorescent labeling, protein samples were buffer exchanged to 10 mM sodium phosphate, pH 8.3, using Micro Bio-Spin 6 columns (Bio-Rad). NHS-ester dyes were prepared immediately before use and mixed with protein at a dye-to-protein molar ratio of 2:1. Tau was labeled using Atto655 NHS-ester (Sigma-Aldrich, 76245), dissolved in acetonitrile, whereas BRICHOS was labeled using Atto390 NHS-ester (Sigma-Aldrich, 68321), dissolved in DMSO. Labeling reactions were incubated at room temperature for 30 min. Unbound dye was removed using Micro Bio-Spin 6 columns (Bio-Rad). Absorbance at 280 nm and at the dye-specific absorption maximum was measured and used to calculate the degree of labeling (DOL). The labeling efficiency was approximately 80% for tau and 30% for BRICHOS.

Tubulin purified from porcine brain (Cytoskeleton, Inc., T240) was dissolved in MilliQ water and buffer exchanged into 20 mM sodium phosphate, pH 8.0, using Micro Bio-Spin 6 columns prior to labeling. Atto488 NHS-ester was dissolved in acetonitrile and added to tubulin at a 1:1 dye-to-protein molar ratio. Following incubation at room temperature for 30 min, excess dye was removed by two consecutive Bio-Spin desalting steps into 20 mM ammonium acetate (AmAc), pH 8.3. The labeling efficiency was approximately 88% per tubulin monomer.

**Computational modeling**

Tau protein was divided into nine non-overlapping 49 residue segments. Individual Tau fragments–BRICHOS interactions were modeled using AlphaFold2 using ColabFold v1.5.5 with MMseqs2 [4]. The MSA mode was set to MMseqs2 (UniRef + environmental), pair mode was selected as “unpaired + paired,” and the AlphaFold-multimer-v2 model was employed with three recycling steps. No template information was used, and the highest-ranked complex was subjected to Amber relaxation. The highest-ranked model was selected and tau chain interface predicted TM-scores (ipTM) were extracted and used for comparative analysis. For Tau fragment interactions with α- and β-tubulin, overall interface confidence was quantified using the Tau chain ipTM score, extracted from the chain_iptm values of the AlphaFold-multimer summary_confidences output, which reflects the interaction of Tau with all other chains in the complex.

Interactions of BRICHOS and tubulin with Tau residues 197–274 were modeled using the AlphaFold3 web server [5]. Predicted complexes were analyzed to identify the relative binding sites of BRICHOS and tubulin on Tau, and per-residue confidence scores (pLDDT) were used to assess local model reliability.

**Mass Photometry**

All mass photometry experiments were carried out at room temperature using the Refeyn TwoMP instrument (Refeyn) in the normal measurement mode with regular image size.

All proteins were prediluted in 25 mM AmAc pH8 (Tau: 500 nM; Tubulin: 250 nM; BRICHOS: 1 µM). These predilutions were then used to incubate the proteins in the desired buffers at 2x the final concentration for 2 min. After focusing with 10 µL buffer, 10 µL of the incubation mix were used for measurements (Table S1). Data was recorded for 1 min (3000 frames) with the AcquireMP (2024 R1.1) software and histograms for each measurement were created in DiscoverMP (v2024 R1) with β-amylase (56, 112 and 224 kDa) or bovine serum albumin (66 and 132 kDa) in 25 mM AmAc as mass calibrants.

**Native mass spectrometry.**

Protein stocks (200 µM) were buffer exchanged into 25 mM AmAc pH8 (ZebaSpin) and then further diluted with water or 100 mM AmAc to get to the final protein (20 µM) and AmAc concentration. Samples were loaded into nESI capillaries (Thermo Fisher Scientific Inc., MA, USA). Mass spectra were recorded in positive ionization mode on a Waters Synapt G1 traveling-wave IM mass spectrometer (MS Vision, The Netherlands) with a capillary voltage of 1.5 kV and the sample cone at 50 V. Source temperature was 30 ºC and trap and transfer collision energy were 10 V. The trap gas was argon at 5 mL/min and IMS gas was nitrogen at 30 mL/min flow rate. For ion mobility measurements the IMS wave height was set to 13 V and the IMS wave velocity to 360 m/s. Data were analyzed using MassLynx 4.1 (Waters, UK).

**Tubulin polymerization assay**

For tubulin polymerization assays, Tau, BRICHOS, and tubulin were buffer exchanged into 25 mM AmAc, pH 8.0, using Micro Bio-Spin desalting columns. Final assay concentrations were 20 µM Tau (98% unlabeled, 2% fluorescently labeled), 5 µM tubulin (95% unlabeled, 5% fluorescently labeled), and 2 µM BRICHOS. Magnesium acetate (MgAc) was added to a final concentration of 250 µM, and polymerization was initiated by addition of GTP (dissolved in Milli-Q water) to a final concentration of 1 mM. Samples were prepared in black half-area, non-binding 96-well polystyrene microplates with a transparent bottom (Corning, CLS3881) and incubated at 37 °C. Fluorescence microscopy images were acquired (at room temperature) immediately after GTP addition (t = 0 min) and after 30 and 90 min of incubation. Control samples were prepared in the absence of BRICHOS.

**Microscopy**

Fluorescence microscopy images were acquired using a Zeiss LSM700 inverted point-scanning confocal microscope (Carl Zeiss, Germany) equipped with a 40× water-immersion objective (NA 1.2). Samples were prepared in non-binding black half-area 96-well polystyrene microplates with a transparent bottom (Corning, CLS3881). Imaging was performed at room temperature. Image acquisition was performed using Zeiss ZEN Blue software (Zeiss), and image analysis was carried out using Fiji.

### MD simulations

The initial structure of the Tau peptide derivative (residues 217-235) was generated in an extended conformation using the TLEAP module of AMBER 24. The amino- and carboxy-terminus were acetylated and amidated respectively. It was subjected to a short Molecular Dynamics (MD) simulation in vacuum for 200 ps, and placed in the center of a truncated octahedral box such that the minimum distance between any peptide atom and the box boundaries was at-least 12 Å. The force-field parameters of ammonium and acetate ions were derived using the GAFF2[6] force-field and bcc charges through the ANTECHAMBER module of AMBER 24. The net charge of ammonium and acetate was set to “+1e”, and “-1e” respectively. The appropriate number of ammonium and acetate ions equivalent to 30 mM (n=2), 100 mM (n=8) and 200 mM (n=16) salt concentrations was computed based on the volume of the primary simulation box (133352 Å$^3$). Packmol[7] was used to generate the three initial distributions of the ammonium and acetate ions within a sphere of radius 20 Å from the centre-of-mass of the peptide. Solvation was performed with the OPC[8] water model and the net charge of the peptide “+4e” was neutralized by adding four acetate counterions. MD simulations were performed using the PMEMD module of AMBER 24 with ff19SB[9] force-field parameters. All the three systems were initially relaxed through energy minimization with steepest descent and conjugate gradient algorithms. They were then heated to 300 K over 30 ps and equilibrated for 200 ps under NVT and NPT ensembles respectively. Production dynamics was run with NPT conditions for 1 µs each. The regulation of the simulation temperature (300 K) and pressure (1 atm), computation of electrostatic interactions, treatment of hydrogen containing bonds and integration of the equation of motion were implemented as described in Lama et al.[10]

### Sequences

| Protein | Sequence |
|---|---|
| hTau441 | MAEPRQEFEVMEDHAGTYGLGDRKDQGGYTMHQDQEGDTDAGLK<br>ESPLQTPTEDGSEEPGSETSDAKSTPTAEDVTAPLVDEGAPGKQAA<br>AQPHTEIPEGTTAEEAGIGDTPSLEDEAAGHVTQARMVSKSKDGTG<br>SDDKKAKGADGKTKIATPRGAAPPGQKGQANATRIPAKTPPAPKTPP<br>SSGEPPKSGDRSGYSSPGSPGTPGSRSRTPSLPTPPTREPKKVAVV<br>RTPPKSPSSAKSRLQTAPVPMPDLKNVKSKIGSTENLKHQPGGGKV<br>QIINKKLDLSNVQSKCGSKDNIKHVPGGGSVQIVYKPVDLSKVTSKC<br>GSLGNIHHKPGGGQVEVKSEKLDFKDRVQSKIGSLDNITHVPGGGN<br>KKIETHKLTFRENAKAKTDHGAEIVYKSPVVSGDTSPRHLSNVSSTG<br>SIDMVDSPQLATLADEVSASLAKQGL |
| Bri2<br>BRICHOS | QTIEENIKIFEEEEVEFISVPVPEFADSDPANIVHDFNKKLTAYLDLNLD<br>KCYVIPLNTSIVMPPRNLLELLINIKAGTYLPQSYLIHEHMVITDRIENID |

| | |
|---|---|
| (R221E) | HLGFFIYELCHDKETYKL |
| proSP-C BRICHOS (T187R) | GSGMKETAAAKFERQHMDSPDLGTDDDDKAMAHMSQKHTEMVLE MSIGAPEAQQRLALSEHLVTTATFSIGSTGLVVYDYQQLLIAYKPAPG TCCYIMKIAPESIPSLEALTRKVHNFQMECSLQAKPAVPTSKLGQAEG RDAGSAPSGGDPAFLGMAVSRLCGEVPLYYI |
| α-tubulin | MRECISIHVGQAGVQIGNACWELYCLEHGIQPDGQMPSDKTIGGGD DSFNTFFSETGAGKHVPRAVFVDLEPTVIDEVRTGTYRQLFHPEQLIT GKEDAANNYARGHYTIGKEIIDLVLDRIRKLADQCTGLQGFSVFHSFG GGTGSGFTSLLMERLSVDYGKKSKLEFSIYPAPQVSTAVVEPYNSILT THTTLEHSDCAFMVDNEAIYDICRRNLDIERPTYTNLNRLIGQIVSSIT ASLRFDGALNVDLTEFQTNLVPYPRGHFPLATYAPVISAEKAYHEQL SVAEITNACFEPANQMVKCDPRHGKYMACCLLYRGDVVPKDVNAAI ATIKTKRTIQFVDWCPTGFKVGINYEPPTVVPGGDLAKVQRAVCMLS NTTAIAEAWARLDHKFDLMYAKRAFVHWYVGEGMEEGEFSEARED MAALEKDYEEVGVDSV |
| β-tubulin | MREIVHIQAGQCGNQIGAKFWEVISDEHGIDPTGSYHGDSDLQLERI NVYYNEAAGNKYVPRAILVDLEPGTMDSVRSGPFGQIFRPDNFVFG QSGAGNNWAKGHYTEGAELVDSVLDVVRKESESCDCLQGFQLTHS LGGGTGSGMGTLLISKIREEYPDRIMNTFSVVPSPKVSDTVVEPYNA TLSVHQLVENTDETYCIDNEALYDICFRTLKLTTPTYGDLNHLVSATM SGVTTCLRFPGQLNADLRKLAVNMVPFPRLHFFMPGFAPLTSRGSQ QYRALTVPELTQQMFDAKNMMAACDPRHGRYLTVAAVFRGRMSMK EVDEQMLNVQNKNSSYFVEWIPNNVKTAVCDIPPRGLKMSATFIGNS TAIQELFKRISEQFTAMFRRKAFLHWYTGEGMDEMEFTEAESNMND LVSEYQQYQD |

**Table S1.** Protein concentrations used in MP experiments

| Protein | Final concentration |
|---|---|
| Tau | 20 nM |
| Tubulin | 5 nM |
| BRICHOS | 200 nM |
| Tau<br>+ Tubulin | 10 nM<br>+ 5 nM |
| Tau<br>+ BRICHOS | 20 nM<br>+ 200 nM |
| Tubulin<br>+ BRICHOS | 5 nM<br>+ 200 nM |
| Tau<br>+ Tubulin<br>+ BRICHOS | 10 nM<br>+ 5 nM<br>+ 200 nM |

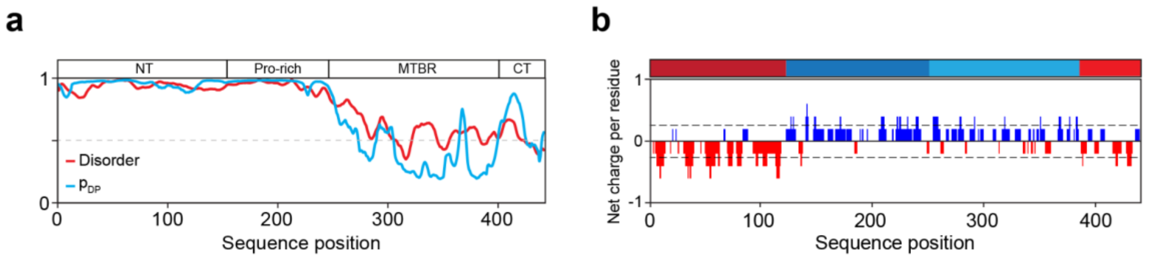


*Figure S1 (a) Overall architecture of tau and predicted disorder (red; IUPred3 webserver[11]) and predicted probability to undergo LLPS (blue; FuzDrop webserver[12]) (b) Net charge per residue distribution of tau in solution with negative charges in red and positive charges in blue (CIDER webserver[13]).*

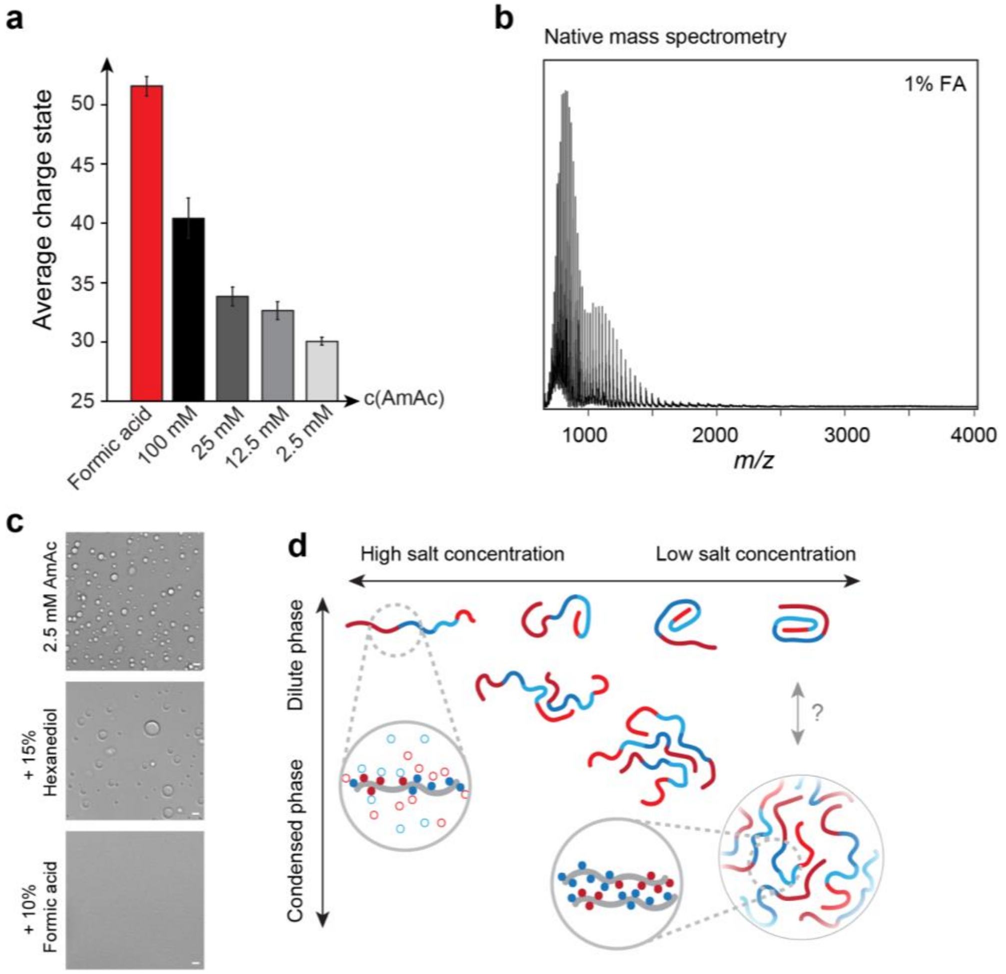


*Figure S2 (a) Average charge states calculated from nMS spectra shown in Figure 2b and Figure S2a. The mean and standard deviation was calculated from three independent measurements.*

(b) Native mass spectrum of tau in 100 mM ammonium acetate (AmAc) pH 8 with 1% (v/v) formic acid. (c) Widefield microscopy of 20 µM tau in 2.5 mM AmAc pH 8 with 15% (v/v) 1,6-hexanediol or 10% (v/v) formic acid added. Scale bars, 5 µm. (d) Schematic overview of the salt-dependent tau LLPS mechanism. Decreasing salt concentrations reduce charge shielding and allow for intra- and intermolecular electrostatic interactions that drive tau LLPS and assembly into condensates. A population of compact tau monomers in the dilute phase is in equilibrium with the condensed phase.

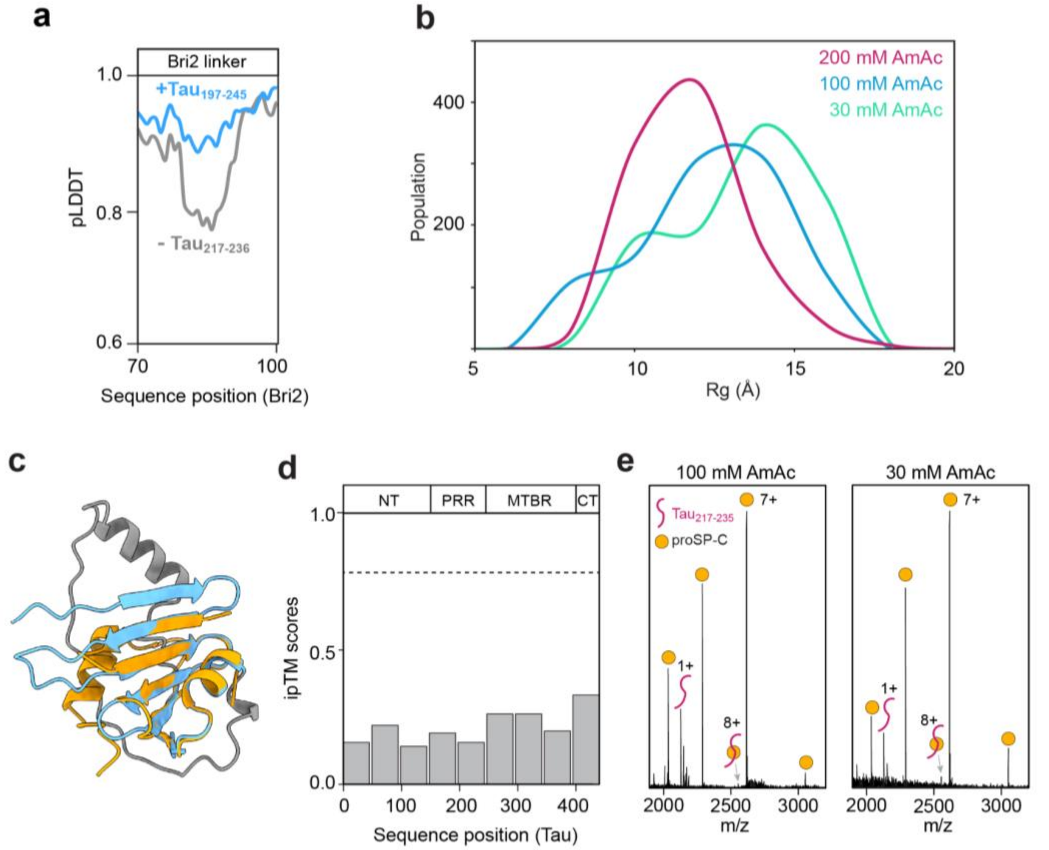


Figure S3. (a) pLDDT score for the Bri2 BRICHOS linker (residue 70-100) (gray). Bri2 linker pLDDT score increases upon tau$_{217\text{-}235}$ binding (blue). Predicted with AlphaFold2. (b) Radius of gyration (Rg) of tau$_{217\text{–}235}$ at decreasing AmAc concentrations, extracted from molecular dynamics simulations. The peptide adopts a more extended conformation under LLPS promoting conditions. (c) Overlay of the Alphafold prediction for the Bri2 BRICHOS domain and the crystal structure of proSP-C BRICHOS (PDB ID: 2YAD) with the conserved core highlighted in blue (Bri2) and orange (proSP-C). (d) ipTM scores for proSP-C BRICHOS binding to 49 residue peptides of tau, predicted with AlphaFold2. No high confidence interactions were predicted (ipTM > 0.8). (e) Native mass spectra of 25 µM tau$_{217\text{-}235}$ (red) with 5 µM proSP-C BRICHOS (orange) in 100 mM and 30 mM AmAc pH 8. Almost no interaction was detected at high or low salt concentration.

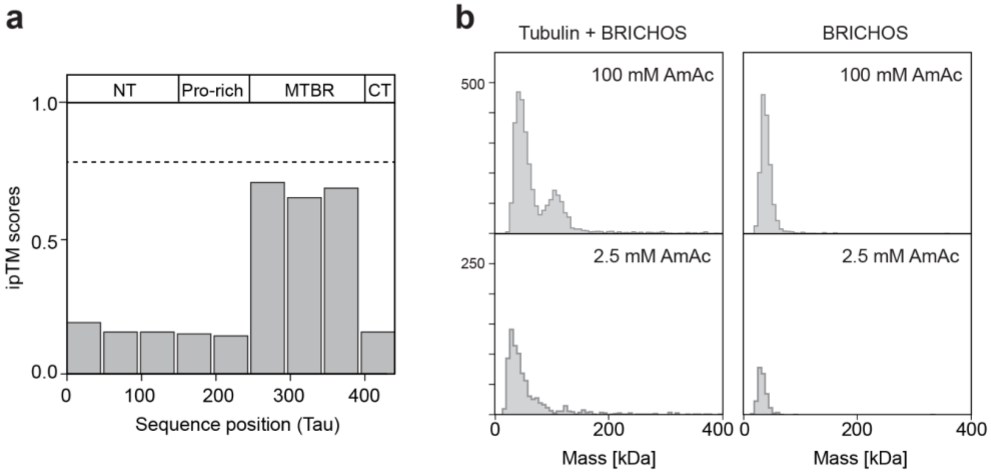


Figure S4. (a) ipTM scores for complexes between 50-residue segments of tau and α/β-tubulin accurately predict preferential interactions of tubulin with the tau MTBR. (b) MP histograms of tubulin (5 nM) with BRICHOS (200 nM) and BRICHOS (200 nM) alone under LLPS and non-LLPS conditions.

### Supplementary References

(1) Danis, C.; Despres, C.; Bessa, L. M.; Malki, I.; Merzougui, H.; Huvent, I.; Qi, H.; Lippens, G.; Cantrelle, F.-X.; Schneider, R.; Hanoulle, X.; Smet-Nocca, C.; Landrieu, I. Nuclear Magnetic Resonance Spectroscopy for the Identification of Multiple Phosphorylations of Intrinsically Disordered Proteins. *J. Vis. Exp. JoVE* **2016**, No. 118, e55001. https://doi.org/10.3791/55001.

(2) Chen, G.; Andrade-Talavera, Y.; Tambaro, S.; Leppert, A.; Nilsson, H. E.; Zhong, X.; Landreh, M.; Nilsson, P.; Hebert, H.; Biverstål, H.; Fisahn, A.; Abelein, A.; Johansson, J. Augmentation of Bri2 Molecular Chaperone Activity against Amyloid-β Reduces Neurotoxicity in Mouse Hippocampus in Vitro. *Commun. Biol.* **2020**, *3* (1), 32. https://doi.org/10.1038/s42003-020-0757-z.

(3) Leppert, A.; Tiiman, A.; Kronqvist, N.; Landreh, M.; Abelein, A.; Vukojević, V.; Johansson, J. Smallest Secondary Nucleation Competent Aβ Aggregates Probed by an ATP-Independent Molecular Chaperone Domain. *Biochemistry* **2021**, *60* (9), 678–688. https://doi.org/10.1021/acs.biochem.1c00003.

(4) Mirdita, M.; Schütze, K.; Moriwaki, Y.; Heo, L.; Ovchinnikov, S.; Steinegger, M. ColabFold: Making Protein Folding Accessible to All. *Nat. Methods* **2022**, *19* (6), 679–682. https://doi.org/10.1038/s41592-022-01488-1.

(5) Abramson, J.; Adler, J.; Dunger, J.; Evans, R.; Green, T.; Pritzel, A.; Ronneberger, O.; Willmore, L.; Ballard, A. J.; Bambrick, J.; Bodenstein, S. W.; Evans, D. A.; Hung, C.-C.; O'Neill, M.; Reiman, D.; Tunyasuvunakool, K.; Wu, Z.; Žemgulytė, A.; Arvaniti, E.; Beattie, C.; Bertolli, O.; Bridgland, A.; Cherepanov, A.; Congreve, M.; Cowen-Rivers, A. I.; Cowie, A.; Figurnov, M.; Fuchs, F. B.; Gladman, H.; Jain, R.; Khan, Y. A.; Low, C. M. R.; Perlin, K.; Potapenko, A.; Savy, P.; Singh, S.; Stecula, A.; Thillaisundaram, A.; Tong, C.; Yakneen, S.; Zhong, E. D.; Zielinski, M.; Žídek, A.; Bapst, V.; Kohli, P.; Jaderberg, M.; Hassabis, D.; Jumper, J. M.

Accurate Structure Prediction of Biomolecular Interactions with AlphaFold 3. *Nature* **2024**, *630* (8016), 493–500. https://doi.org/10.1038/s41586-024-07487-w.
(6) Wang, J.; Wolf, R. M.; Caldwell, J. W.; Kollman, P. A.; Case, D. A. Development and Testing of a General Amber Force Field. *J. Comput. Chem.* **2004**, *25* (9), 1157–1174. https://doi.org/10.1002/jcc.20035.
(7) Martínez, L.; Andrade, R.; Birgin, E. G.; Martínez, J. M. PACKMOL: A package for building initial configurations for molecular dynamics simulations. *J. Comput. Chem.* **2009**, *30* (13), 2157–2164. https://doi.org/10.1002/jcc.21224.
(8) Izadi, S.; Anandakrishnan, R.; Onufriev, A. V. Building Water Models: A Different Approach. *J. Phys. Chem. Lett.* **2014**, *5* (21), 3863–3871. https://doi.org/10.1021/jz501780a.
(9) Tian, C.; Kasavajhala, K.; Belfon, K. A. A.; Raguette, L.; Huang, H.; Migues, A. N.; Bickel, J.; Wang, Y.; Pincay, J.; Wu, Q.; Simmerling, C. ff19SB: Amino-Acid-Specific Protein Backbone Parameters Trained against Quantum Mechanics Energy Surfaces in Solution. *J. Chem. Theory Comput.* **2020**, *16* (1), 528–552. https://doi.org/10.1021/acs.jctc.9b00591.
(10) Lama, D.; Vosselman, T.; Sahin, C.; Liaño-Pons, J.; Cerrato, C. P.; Nilsson, L.; Teilum, K.; Lane, D. P.; Landreh, M.; Arsenian Henriksson, M. A Druggable Conformational Switch in the C-MYC Transactivation Domain. *Nat. Commun.* **2024**, *15*, 1865. https://doi.org/10.1038/s41467-024-45826-7.
(11) Erdős, G.; Pajkos, M.; Dosztányi, Z. IUPred3: Prediction of Protein Disorder Enhanced with Unambiguous Experimental Annotation and Visualization of Evolutionary Conservation. *Nucleic Acids Res.* **2021**, *49* (W1), W297–W303. https://doi.org/10.1093/nar/gkab408.
(12) Hatos, A.; Tosatto, S. C. E.; Vendruscolo, M.; Fuxreiter, M. FuzDrop on AlphaFold: Visualizing the Sequence-Dependent Propensity of Liquid–Liquid Phase Separation and Aggregation of Proteins. *Nucleic Acids Res.* **2022**, *50* (W1), W337–W344. https://doi.org/10.1093/nar/gkac386.
(13) Holehouse, A. S.; Das, R. K.; Ahad, J. N.; Richardson, M. O. G.; Pappu, R. V. CIDER: Resources to Analyze Sequence-Ensemble Relationships of Intrinsically Disordered Proteins. *Biophys. J.* **2017**, *112* (1), 16–21. https://doi.org/10.1016/j.bpj.2016.11.3200.
